# Bayesian Inference of Dependent Population Dynamics in Coalescent Models

**DOI:** 10.1101/2022.05.22.492976

**Authors:** Lorenzo Cappello, Jaehee Kim, Julia Palacios

## Abstract

The coalescent is a powerful statistical framework that allows us to infer past population dynamics leveraging the ancestral relationships reconstructed from sampled molecular sequence data. In many biomedical applications, such as in the study of infectious diseases, cell development, and tumorgenesis, several distinct populations share evolutionary history and therefore become dependent. The inference of such dependence is a highly important, yet a challenging problem. With advances in sequencing technologies, we are well positioned to exploit the wealth of high-resolution biological data for tackling this problem. Here, we present a novel probabilistic model that relies on jointly distributed Markov random fields. We use this model to estimate past population dynamics of dependent populations and to quantify their degree of dependence. An essential feature of our approach is the ability to track the time-varying association between the populations while making minimal assumptions on their functional shapes via Markov random field priors. We provide nonparametric estimators, extensions of our base model that integrate multiple data sources, and fast scalable inference algorithms. We test our method using simulated data under various dependent population histories and demonstrate the utility of our model in shedding light on evolutionary histories of different variants of SARS-CoV-2.

## 1 Introduction

Bayesian inference of past population sizes from genetic data is an important task in molecular epidemiology of infectious diseases and other biomedical disciplines (Volz et al., 2009; Bouckaert et al., 2019; Stadler et al., 2021). While many computational and methodological advances have been developed in the last 20 years, there are still many challenges in using these models on real data applications (See Cappello et al. (2021) for a recent review).

An important limitation of current methods is their lack of flexibility in modeling dependent populations. Current models typically assume either single population size dynamics or a structured population undergoing migration. In particular, when modeling structured populations, these models resort to simplistic assumptions on the population size dynamics in order to gain computational tractability and parameter identifiability (Kühnert et al., 2016; Müller et al., 2017). However, often there are situations where the population is neither one large unit nor completely divided into isolated subpopulations. Different subpopulations may share the same environment and ancestry, and therefore their population dynamics are dependent. For example, in the study of infectious diseases, a new variant can emerge and become dominant over the preexisting variants in the population. In this case, all variants share the same environment and local non-pharmaceutical interventions. In tumorgenesis, the cancer cell population within an individual undergoes clonal expansions in the tumor microenvironment, often resulting in a mixture of genotypically and phenotypically heterogenous cell subpopulations. Identifying and quantifying such expansions is crucial for timely detection and personalized oncology for cancer (Caswell-Jin et al., 2021). Despite its importance, to the best of our knowledge, no realistic methods exist for jointly modeling and studying the evolutionary trajectories of dependent populations.

In this work, we propose a nonparametric method for inferring past population dynamics of dependent populations and for estimating the relative difference in their evolutionary trajectories over time. The proposed method bypasses the problems inherent in modeling complex dependent population dynamics by *a priori* modeling the dependency of population sizes via Gaussian Markov random fields. Our approach incorporates other types of data informative of the parameter of interest such as temporal sampling information of molecular sequences.

Our contributions can be summarized as follows.

- We propose a nonparametric Bayesian framework for inferring dependent population dynamics from genetic data in the coalescent framework. The model makes minimal assumptions on the functional form of the population trajectories and their dependency. Despite its flexibility, we prove that model parameters are identifiable.
- We extend our framework to jointly model the ancestral and sampling processes incorporating sampling times as an additional source of information. We empirically validate the performance of our methods and show the ability of the sampling-aware methods in reducing bias and improving estimation.
- We demonstrate the utility of our methods in providing new insights into the evolutionary dynamics of SARS-CoV-2 novel variants.

## 2 Background

### 2.1 Coalescent Model

The coalescent (Kingman, 1982) is a popular prior model on genealogies. The genealogies are timed and rooted binary trees that represent the ancestry of a sample of *n* individuals from a population. We assume that (*n_ℓ_*)_*ℓ*=1:m_ samples are sequentially collected at *m* sampling times denoted by **s** = (*s_ℓ_*)*_ℓ=1:m_*, with *s*_1_ =0, *S*_*j*-1_ < *S_j_* for *j* = 2,…, *m*, and 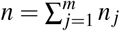 is the total number of samples. In the genealogy, pairs of lineages merge backwards in time into a common ancestor at coalescent times denoted by **t** = (*t_n_*,…, *t*_2_) (Figure 1). The rate at which pairs of lineages coalesce depends on the number of lineages and the effective population size (EPS) denoted by (*N_e_*(*t*))_t≥0_:=*N_e_*. Under this model, the density of a genealogy **g** is:

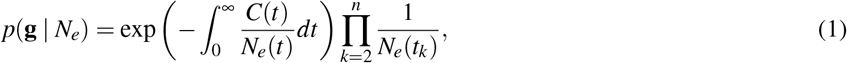

where 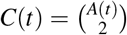 is the coalescent factor—a combinatorial factor of the number of extant lineages 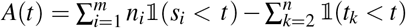. The EPS is generally interpreted as a relative measure of genetic diversity (Wakeley & Sargsyan, 2009).

**Figure 1.**
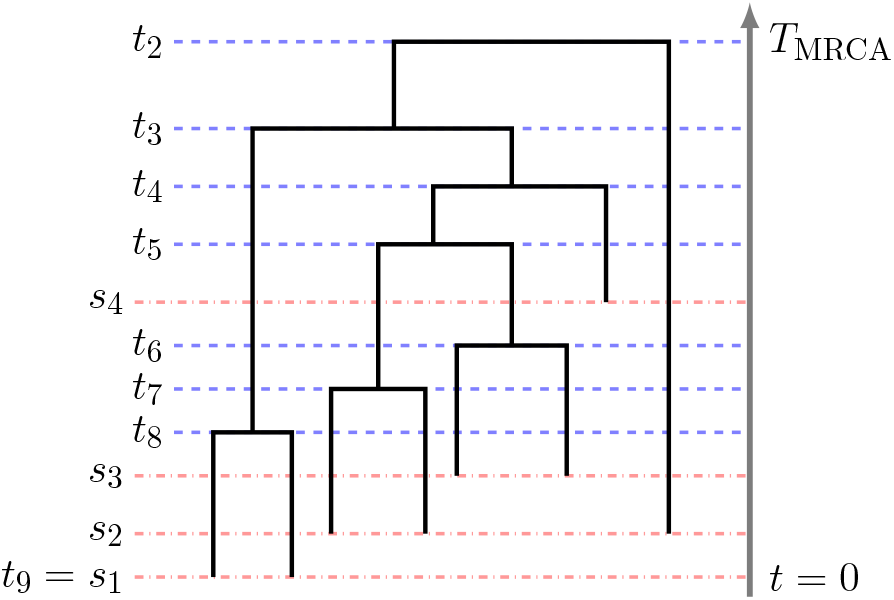
Example of a Genealogy with Sequential Sampling. *s*_1_,…, *s*_4_ and *t*_8_,…, *t*_2_ indicate sampling times (red dotted lines) and coalescent times (blue dotted lines), respectively. The time increases backwards in time starting with *s*_1_ = *t*_9_ = 0 as the present time. *T_MRCA_* indicates the time to the most recent common ancestor at the root.

### 2.2 Bayesian Inference of EPS

We start by assuming that a given genealogy is available to us. Bayesian inference of the EPS then targets

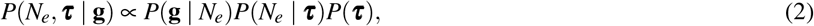

where *P*(*N_e_* | ***τ***) is a prior distribution on *N_e_* that depends on a vector of hyperparameters ***τ***.

A common choice for a prior on *N_e_* is to assume an underlying finite-dimensional parametric structure. In particular, this choice makes the calculation of the integral in Eq. (1) computationally tractable. A popular strategy is to use a regular grid of *M* + 1 points (*k_i_*)_1:*M*+1_ and assume that *N_e_* is well approximated by

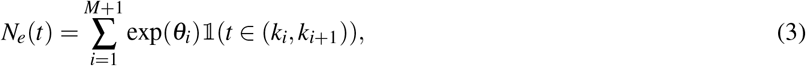

a piece-wise constant function with *M* change points. In Eq. (3), we model *N_e_* in log-scale, however this is not strictly necessary. Many approaches place a Gaussian Markov random field (GMRF) prior on ***θ***, for example Gill et al. (2013); Volz & Didelot(2018); Faulkner et al. (2020). An alternative to the piece-wise constant assumption is to use Gaussian process priors. However, the posterior distribution becomes doubly intractable because the likelihood function depends on an infinite-dimensional integral over Gaussian processes. In Palacios & Minin (2013), the authors augmented the posterior distribution with auxiliary variables via thinning of Poisson processes (Adams et al., 2009) in order to gain tractability.

In applications, the use of GMRFs has shown a numerical performance comparable to GPs. We conjecture that the reason of this comparable performance is the flexibility in the choice of the number of parameters *M*. This is the case because neither the grid cell boundaries (*k_i_*)_1:*M*+1_ nor *M* depend on the data, **g** = (*g*, **t**, **s**, **n**) or the model parameters *N_e_*(*t*). We refer to Faulkner et al. (2020) for guidance on the choice of *M*.

There are exact and approximate algorithms to sample from Eq. (2). Standard software packages (Suchard et al., 2018; Bouckaert et al., 2019) employ Markov chain Monte Carlo (MCMC) methods with carefully designed transition kernels. Recent algorithmic advances employ Hamiltonian Monte Carlo (HMC).Faulkner et al. (2020) implemented an algorithm to sample from Eq. (2) under several Markov random fields priors on STAN (Carpenter et al., 2017), and Lan et al. (2015) employed split HMC (Shahbaba et al., 2014). Among the approximate methods, inference can be efficiently done with Integrated Nested Laplace Approximation (INLA) (Rue et al., 2009).Palacios & Minin (2012) use INLA under a GMRF prior on *N_e_*, showing that the approximate posterior is remarkably similar to that obtained by MCMC-based algorithms.

### 2.3 Preferential Sampling

The standard coalescent model implicitly assumes the sampling times are either fixed or functionally independent of the underlying population dynamics. However, in many applications, such as in infectious diseases, the sampling frequency is often highly correlated with EPS: more samples are sequenced when the EPS is larger. This situation, known as preferential sampling in spatial statistics (Diggle et al., 2010), allows us to model sampling frequency information in order to improve inference about EPS, reducing estimation bias and improving the accuracy of model parameter inference (Volz & Frost, 2014; Karcher et al., 2016, 2020; Parag et al., 2020; Cappello & Palacios, 2021).

The probabilistic dependency of sampling time distribution on population dynamics can be modeled as an inhomogeneous Poisson point process (iPPP) with a rate *λ*(*t*) that depends on EPS. We adopt the adaptive preferential sampling framework (Cappello & Palacios, 2021) that employs a flexible approach for modeling the time-varying dependency between *N_e_*(*t*) and *λ*(*t*): *λ*(*t*) = *ζ*(*t*)*N_e_*(*t*), where both *ζ*(*t*) and *N_e_*(*t*) are unknown continuous functions with GMRF priors.

## 3 Methods

Here, we propose a flexible and scalable framework for modeling the genealogies of two samples (not necessarily of the same size) from two populations with dependent population dynamics. The goals are (i) to estimate the EPSs of the two populations, (ii) to account for the dependency between them, (iii) and to quantify estimation uncertainty.

### 3.1 Coalescent for Dependent Evolutionary Histories

Let ***g**^A^* = (*g^A^*, **t**^A^, **s**^A^, **n^A^**) and **g^B^** = (*g^B^*, **t**^*B*^, **s**^*B*^, **n**^*B*^) be the genealogies of samples collected from populations *A* and *B* respectively, and let 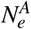 and 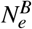 denote their corresponding EPSs. Here, (**s**^*A*^, **s**^*B*^) are vectors of sampling times, and (**n**^*A*^, **n**^*B*^) are number of samples collected. Standard coalescent-based inference methodologies approximate posterior distributions 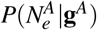 and 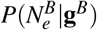 ignoring any association between the underlying population processes of the two populations.

We model the association between the two population size trajectories, which can change over time, with a time-varying parameter linking 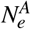 and 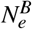:

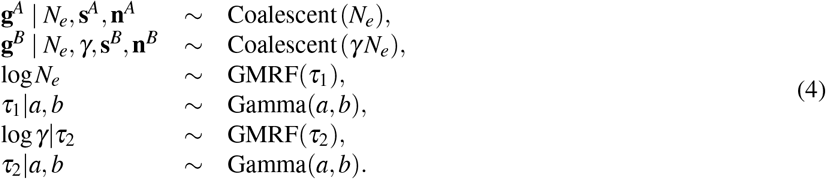

Here, *γ*:= (*γ*(*t*))_*t*≥0_ is the time-varying coefficient that describes how the association between the two population processes changes over time, leading to 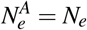 and 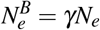. The interpretation of *γ* provides information on the association between two populations. For example, a growing trend in *γ* signals the existence of an association between two EPSs: 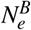 is growing faster than 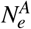. It, however, does not necessarily imply a positive association because, for example, if 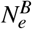 were growing, 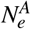 could be either growing at a slower rate than 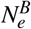 or be decreasing and still have an increasing *γ*.

The number of parameters of the two GMRFs tunes how free the dependence is allowed to vary between the two populations. Throughout the section, we employ GMRF priors on *N_e_* and *γ*; however, the framework is flexible to any prior distribution. It is possible to go fully nonparametric employing a GP (Palacios & Minin, 2013), or a different kind of MRFs, for example, the Horseshoe MRFs (Faulkner & Minin,2018).

Our model in Eq. (4) is “asymmetric”, in the sense that the baseline population EPS is tilted by the time-varying coefficient *γ* to define the EPS of a new population. The choice is motivated by the actual scientific question we are examining, where a new population develops from an existing one. An alternative model is to allow every EPS to be a function of a shared base population size trajectory. This second formulation is more symmetrical, although the parameters lose interpretability and are not identifiable.

We also considered a simpler parametric model suggested in Cappello et al. (2021). Here, the population dependence is modeled through two time-independent parameters *α* and *β*:

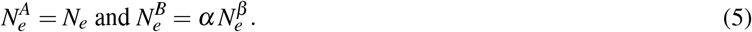

However, the strict parametric dependence enforced in the model increases the risk of model misspecification by not allowing changes in the association of the two population processes. For example, the model support excludes a time shift when 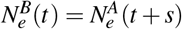. A consequence of this potential model misspecification is biased estimation of 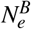, *α*, and *β*; we will provide numerical proofs of this claim in Section 4.

### 3.2 Preferential Sampling and Dependent Evolutionary Histories

A common approach for studying the relative growth between two population dynamics is to model how the sampling frequency of molecular sequences changes over time in the two populations. This is frequently done by fitting logistic growth models to the sampling dates only (Volz et al., 2021; Davies et al., 2021). The probabilistic models discussed in Section 3.1 take a different stance and employ molecular data to reconstruct the genealogies, which in turn are used to infer the population processes jointly. Here, we extend the probabilistic models of Section 3.1, either model Eq. (4) or (5), and model the genealogies jointly with the observed sampling frequencies from both populations as follows:

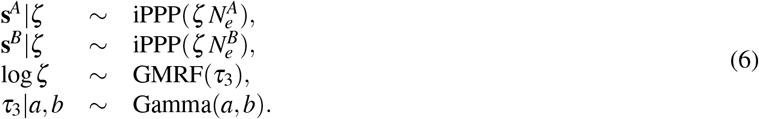

Eq. (6) builds on the preferential sampling framework described in Section 2.3: the sampling process is an iPPP whose rate is a function of both, the EPS and a time-varying parameter *ζ*. Here, the sampling process of population *A* and *B* will have distinct rates, 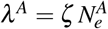 and 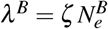. Although we assumed a shared *ζ* function, our implementation considers the possibility of one *ζ* function per population.

### 3.3 Inference

We start describing the inference procedure for the parameters of the model in Eq. (4). We employ the same discretization described in Section 2.2: given a regular grid (*k_i_*)_1:*M*+1_, we assume that *N_e_* is governed by parameters ***θ*** through the map given in Eq. (3); *γ* is governed by parameters **ζ** = (*ζ_i_*)_*i*=1:*M*_’ such that *γ*(*t*) ≈ exp *ξ_i_* for *t* ∈(*k_i_*, *k*_*i*+1_]. Let ***τ*** be the vector of precision hyperparameters of the GMRFs. Based on Palacios & Minin (2012), we use INLA for obtaining marginal posterior medians and marginal 95% Bayesian credible intervals (BCI).

INLA does not approximate the full posterior *P*(***ζ***, ***θ***, ***τ***|**g^*A*^**, **g^*B*^**); rather, it approximates the posterior marginals *P*(***τ***|**g^*A*^**,**g^*B*^**), (*P*(*θ_i_*|**g^*A*^**, **g^*B*^**))_1:*M*_, and (*P*(*ξ_i_*|**g**^*A*^, **g**^*B*^))_1:*M*_. The first step consists in computing

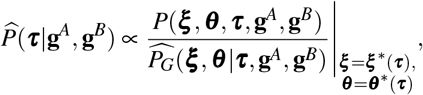

where the denominator is the Gaussian approximation to *P*(***ζ***, ***θ***|***τ***, **g**^*A*^, **g**^*B*^) obtained from a Taylor expansion around its modes ***θ***\*(***τ***) and ***ζ***\*(***τ***) (the first Laplace approximation). The second step approximates the marginal posteriors of *P*(*θ_i_*|**g**^*A*^, **g**^*B*^) and *P*(*ξ_i_*|**g**^*A*^, **g^*B*^**). For example *P*(*θ_i_*,|**g^*A*^**, **g^*B*^**) is approximated by

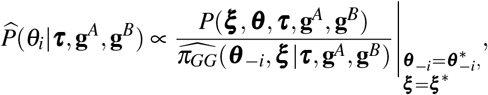

where the denominator is again a Gaussian approximation of the corresponding conditional distribution. Now the Taylor expansion is centered at (**θ**_*-i*_,) = E_G_[**θ**_*-i*_, *ξ*|**τ**,**g**^*A*^,**g**^*B*^], where the expected value is taken w.r.t. 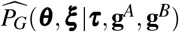. The last step involves integrating out the hyperparameters from 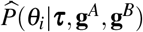. This can be easily accomplished using 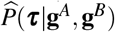 (the nested Laplace approximation step).

### 3.4 Identifiability

Parameter identifiability is an essential property of models used in statistical learning. Roughly speaking, it refers to the theoretical possibility of uniquely estimating a parameter vector if an infinite amount of data is available (Rothenberg, 1971; Bishop, 2006; Watanabe, 2009). Note that this is a property of the generative model, not of the estimator used.

#### Definition 3.1

(Lehmann & Casella (2006)). Let *X* be distributed according to *P_θ_* with *θ* ∈ **Θ**, *θ* is said to be <u>identifiable</u> if there exist no *θ*_1_ ≠ *θ*_2_, with *θ*_1_, *θ*_2_ ∈ **Θ**, for which *P_θ_1__* = *P_θ_2__*.

Definition 1 can be rewritten stating that a parameter is identifiable if the map *θ* ↦ *P_θ_* is one-to-one. A model is nonidentifiable if two or more parameter vectors are observationally equivalent. An example of nonidentifiable parameters arises if *Y* ~ Poisson(*λ*_1_*λ*_2_): for a pair 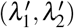, any combination 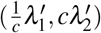 with *c* > 0 will be observationally equivalent. Parameter redundancy is another common violation (Catchpole & Morgan, 1997).

The parameters of models in Eqs. (4) and (5) have a scientific interpretation. The lack of identifiability could hinder the validity of the scientific insights gained from using our methodology. There is a large literature showing that many models in evolutionary biology and ecology are not identifiable (Little et al., 2010; Cappello et al., 2021). Under the assumption that *N_e_* and *γ* are piecewise-constant, *i.e. N_e_* = (*N_e,i_*)_1:*M*_ and *γ* = (*γ_i_*)_1:*M*_, we prove that the models introduced in this section possess this important property.

#### Proposition 3.2

(Identifiability of the nonparametric model, Eq. (4)). *Let g^A^ be distributed as a coalescent with EPS* (*N_e,i_*)_1:*M*_, *and g^B^ as a coalescent with EPS (γ_i_N_e,i_)_1:M_*, *M* ≥ 1, *then the vector (N_e,1_,≥, N_e,M_*, *γ*_1_,…, *γ_M_*) *is identifiable.*

#### Proposition 3.3

(Identifiability of the parametric model, Eq. (5)). *Let g^A^ be distributed as a coalescent with EPS (N_e,i_)_1:M_, and g^B^ as a coalescent with EPS 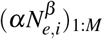, if M ≥ 2, the vector (N_e,1_,≥,N_e,M_, α, β*) *is identifiable.*

Proofs of Propositions 3.2 and 3.3 are included in the Supplementary Material. There are multiple techniques to prove the results stated above; see Ran & Hu (2017) for a review. We employ results in Rothenberg (1971), which consists in showing that the expected Fisher information (EFI) is non-singular.

### 3.5 Extension to multiple populations

Our work is centered on a two-population model because our main application target is the estimation of a relative selective advantage of a newly emerging viral variant over an existing variant. We further assume that the new variant originated from the standing variant. However, the framework can easily be extended to include multiple populations viewing the two-population model as a building block for a general genealogical model with multiple populations. In this generalization, the EPS of a child population is a function of the EPS of its parental population. This gives the two types of hierarchical structures displayed in Figure 2:

- *Nested populations.* Each effective population is a function of its immediate preceding one (Figure 2(A)). The baseline population “A” with EPS 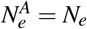 evolves into a second population “B” with EPS 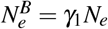, which then in turn evolves into a population “C” with EPS 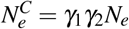.
- *“Radial” populations.* Multiple populations evolve from the baseline population “A” with EPS 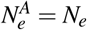 (Figure 2(B)). Then, population “B” will have EPS 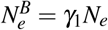, and population “C” 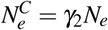.

**Figure 2.**
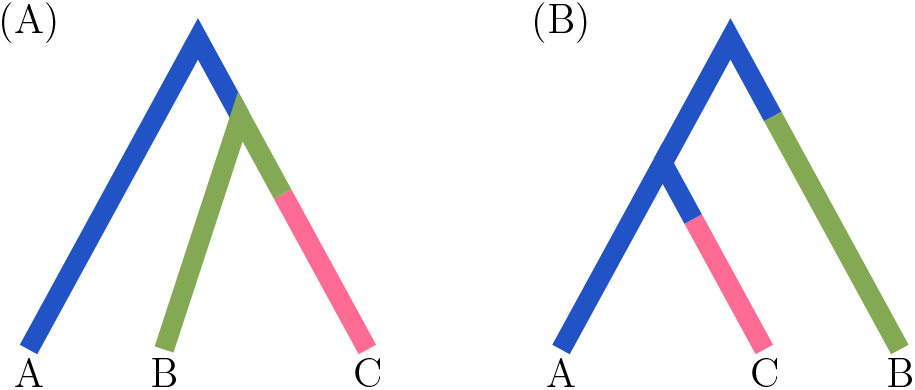
Modeling of Multiple Dependent Populations. The trees represent large genealogies of many sequences from three different populations labeled A, B and C at the tips of the trees. Each lineage represents subtrees of individuals whose rate of coalescence is dictated by the color of the branch. **(A) Nested.** The blue branch indicates coalescent events happen at rate that depends on 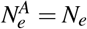 green branch indicates a coalescent rate that depends on 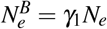 and pink branch indicates a coalescent rate that depends on 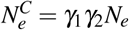. **(B) Radial.** The blue branch indicates a coalescent rate that depends on 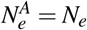, the green branch indicates a coalescent rate that depends on 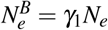 and pink branch indicates a colescent rate that depends on 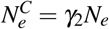.

The hierarchical structure for EPSs of multiple populations can be constructed iteratively as combinations of the two base structures of Figure 2. This construction maintains parameter identifiability. The inference framework for the multiple-populations extension still follows Section 3.3: GMRF priors on the base EPS *N_e_* and the *γ_i_* parameters and the parameter inference with INLA approximation.

We note that, while the above extension can model more general evolutionary scenarios, it still cannot accommodate all possible evolutionary histories. For example, it cannot model ancestral populations that are not observed in the samples.

## 4 Experiments

We show the effectiveness of our methodology by applying it to synthetic and real-world data. In the synthetic data section, we offer evidence of its numerical accuracy. The real-data section illustrates the scientific insights that can be obtained by applying our methodology to SARS-CoV-2 data. The code to reproduce the following studies is available at https://github.com/lorenzocapp/adasel.

### 4.1 Synthetic Data

Figure 3 depicts six pairs 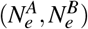 used to simulate data. The trajectories mimic challenging realistic scenarios typically encountered in applications. The scenarios include several types of dependence between 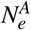 and 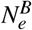, ranging from perfect association (Scenario 6) to no association (Scenario 2; here, our model is misspecified). We simulated 100 datasets per scenario. For a fixed pair of EPSs, we sampled (***s^A^***, ***s^B^***), (***n^A^***, ***n^B^***), and (***t^A^***, ***t^B^***) with *n* = 200. Specifics of 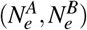 and the data-generating mechanism are in the Supplementary Material.

**Figure 3.**
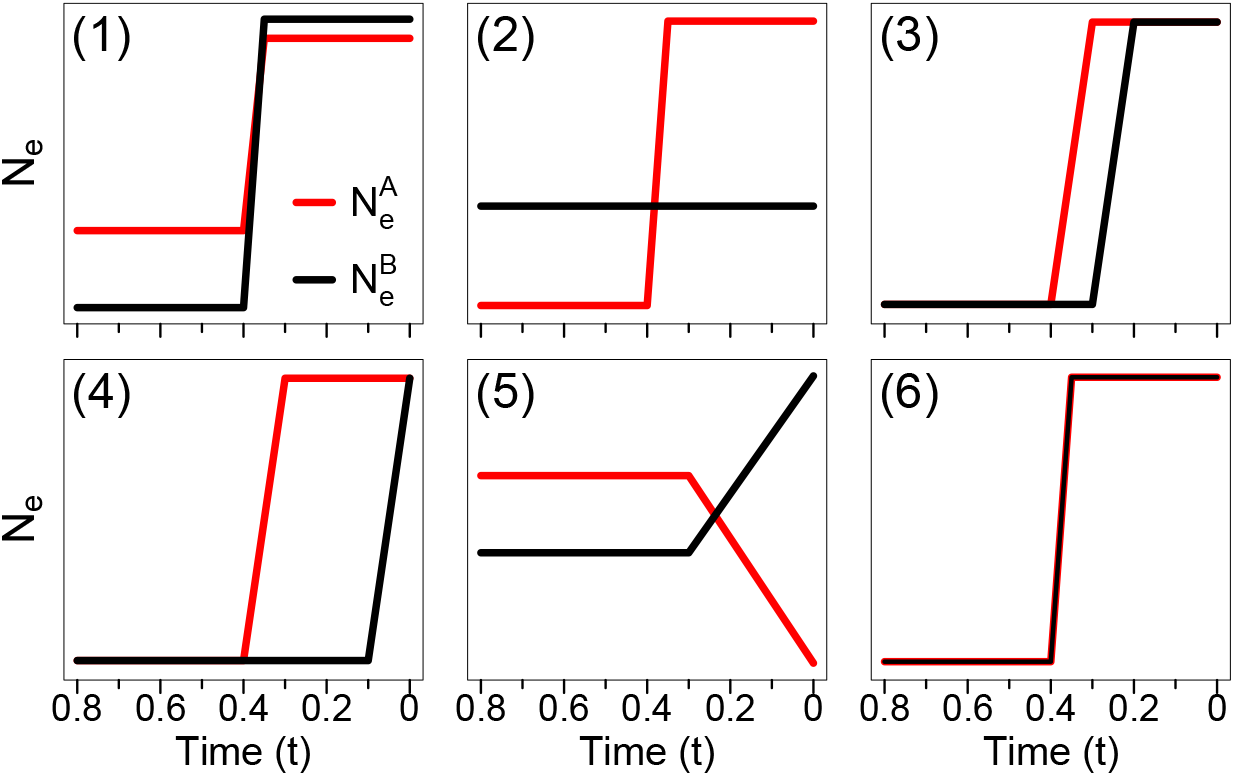
Population Trajectories for Synthetic Data.

We compare our hierarchical approach with and without preferential sampling—referred as “adaPop” and “adaPop+Pref”, respectively—to the parametric method, referred here as “parPop”. We also include a neutral estimator “noPop”, which ignores the association between populations and estimates 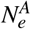 and 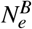. We compare how accurately the four methodologies estimate *γ*, 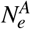 and 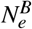. Note that, while parPop and noPop do not approximate *γ*, the posterior *P*(*γ*|***s**^A^s^B^*,**t^*A*^**, **t^*B*^**) can be empirically approximated by taking samples from 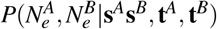 and summarizing 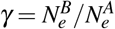. Let *f* be either *γ*, 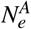 or 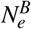. We evaluate the performance of the methods using three metrics (listed below) computed on a regular grid of time points (v_*i*_)_1:*K*_. Here, the grid is defined with *K* = 100 on the interval [0,0.6*T_MRCA_*], where *T_MSCA_* denotes the time to the most recent common ancestor at the root.

- 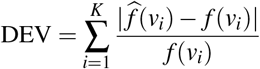, where 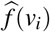 is the posterior median of *f* at time *v_i_*. It is a measure of bias.
- 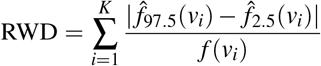, where 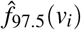 and 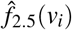 are respectively the 97.5% and 2.5% quantiles of the posterior distribution of *f*(*v_i_*). It describes average width of the credible region.
- 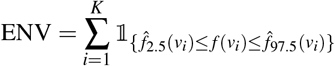. It is a measure of 95% credible intervals coverage.

Table 1 reports the average value of each statistic pooling together all datasets, all scenarios, and all grid points. Hence, each entry should represent the average performance of a method across the variety of challenging scenarios considered. We also average the performance metrics of 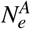 and 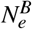. A more granular view of the performance of each method by scenario is given in the Supplementary Material.

**Table 1.**
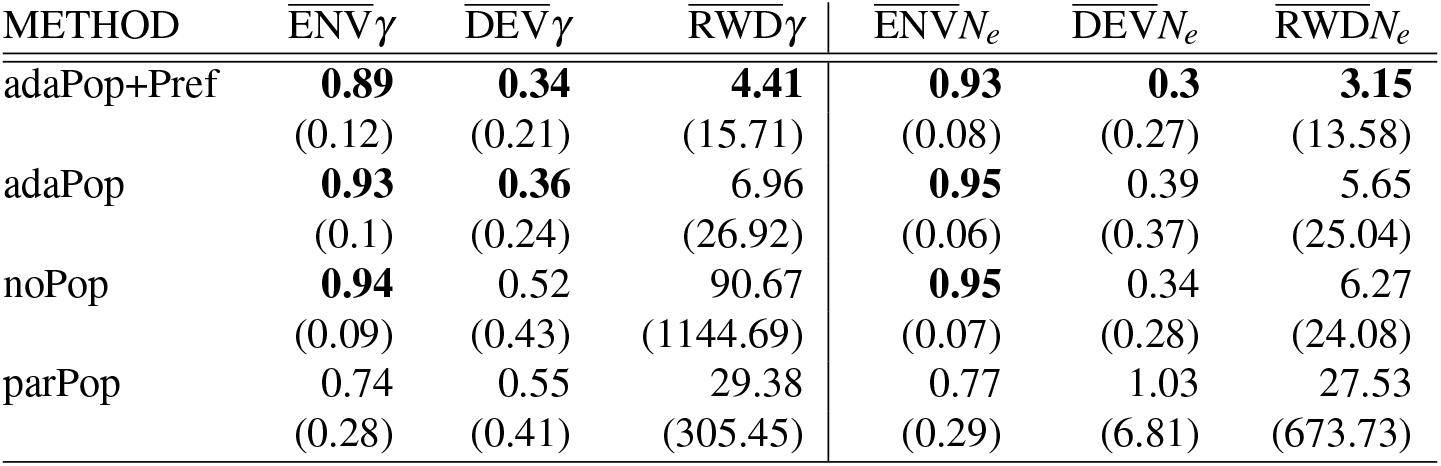
Summary Statistics of Posterior Inference of *γ*, 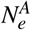, and 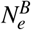. Each entry is computed as the mean of the performance metric considered across all synthetic datasets (six possible scenarios, 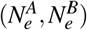, and 100 datasets per scenario). The metrics for 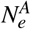 and 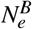 have been also averaged. Numbers in parentheses are the standard deviation of each estimate. The numbers in bold indicate the method(s) with the best performance (and within 10% of the best) for each performance metric: the highest for ENV, the lowest for DEV and RWD.

| METHOD | $\overline{\text{ENV}}\gamma$ | $\overline{\text{DEV}}\gamma$ | $\overline{\text{RWD}}\gamma$ | $\overline{\text{ENV}}N_e$ | $\overline{\text{DEV}}N_e$ | $\overline{\text{RWD}}N_e$ |
| --- | --- | --- | --- | --- | --- | --- |
| adaPop+Pref | **0.89**<br>(0.12) | **0.34**<br>(0.21) | **4.41**<br>(15.71) | **0.93**<br>(0.08) | **0.3**<br>(0.27) | **3.15**<br>(13.58) |
| adaPop | **0.93**<br>(0.1) | **0.36**<br>(0.24) | 6.96<br>(26.92) | **0.95**<br>(0.06) | 0.39<br>(0.37) | 5.65<br>(25.04) |
| noPop | **0.94**<br>(0.09) | 0.52<br>(0.43) | 90.67<br>(1144.69) | **0.95**<br>(0.07) | 0.34<br>(0.28) | 6.27<br>(24.08) |
| parPop | 0.74<br>(0.28) | 0.55<br>(0.41) | 29.38<br>(305.45) | 0.77<br>(0.29) | 1.03<br>(6.81) | 27.53<br>(673.73) |

adaPop+Pref stands out as the best performing method, exhibiting the lowest bias (DEV) and the narrowest credible regions (RWD). The coverage (ENV) is slightly worse than noPop; however, noPop achieves the slightly higher coverage with much wider credible regions. adaPop has a very similar performance to adaPop+Pref.

Sampling times were sampled at uniform and not proportionally to 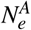 and 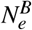. This is the case where preferential sampling is less informative. Remarkably, adaPop+Pref still has narrower credible regions than adaPop. The reason is that the time-varying coefficient *ζ* is able too capture a variety of sampling protocols, including uniform sampling. We interpret this as empirical evidence of the adaptivity of the model.

adaPop’s higher 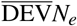 is mostly attributed to poorer performance in estimating 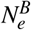 in Scenario 2 (see Supplementary Material). If we exclude Scenario 2, adaPop is superior to noPop and parPop across all metrics. Similarly, parPop is very competitive in the scenarios where the model is correctly specified (Scenarios 1, 2, and 6). The average performance deteriorates due to poorer performance in the remaining scenarios.

Lastly, an essential feature of adaPop and adaPop+Pref is that they are the most stable methodologies. This can be seen from the standard deviations of the statistics in the bracket. The Supplementary Material includes further analyses where the robustness and performance of the models are evaluated.

### 4.2 Real Data

Since its introduction, SARS-CoV-2 has undergone rapid evolution resulting in novel variants, some of which possess transmissibility, pathogenicity, or antigenicity advantages over the preexisting resident variants (Harvey et al., 2021; Cevik et al., 2021). A variant of recent interest is the delta variant (Pango lineage B.1.617.2 and AY lineages (Rambaut et al., 2020)). We compare this variant to other SARS-CoV-2 variants in two countries, South Korea and Italy.

We analyzed high-coverage complete sequences publicly available in GISAID (Shu & McCauley, 2017) collected from South Korea and Italy during 2021-03-01 to 2021-09-30. For each country, we subsampled two sets of 150 sequences: one with the delta variant and the other without the delta variant. We then estimated the maximum credibility clade (MCC) trees—the tree in the posterior sample with the maximum sum of the posterior clade probabilities—of samples from each variant group of each country independently with BEAST2 (Bouckaert et al., 2019). In our study, we set the population of the delta variant sequences as population A, and of the non-delta variant as population B. Here, we discuss the results using the parPop and adaPop+Pref models. The additional results with other methods and further details of the sequence analysis pipeline, together with the inferred MCC trees, appear in the Supplementary Materials.

The orange-shaded heat-maps in Figure 4 depict the number of samples collected over time of the two populations in South Korea and Italy (as represented in our sub-sampled data sets). The observed pattern is consistent in the two countries: delta viral samples predominantly collected in the summer of 2021, while non-delta samples collected in the spring and early summer of 2021. This is consistent with the general observation of rapid spread of the delta variant that has progressively replaced the preexisting non-delta variants (such as alpha) since its introduction (Mlcochova et al., 2021; del Rio et al., 2021).

**Figure 4.**
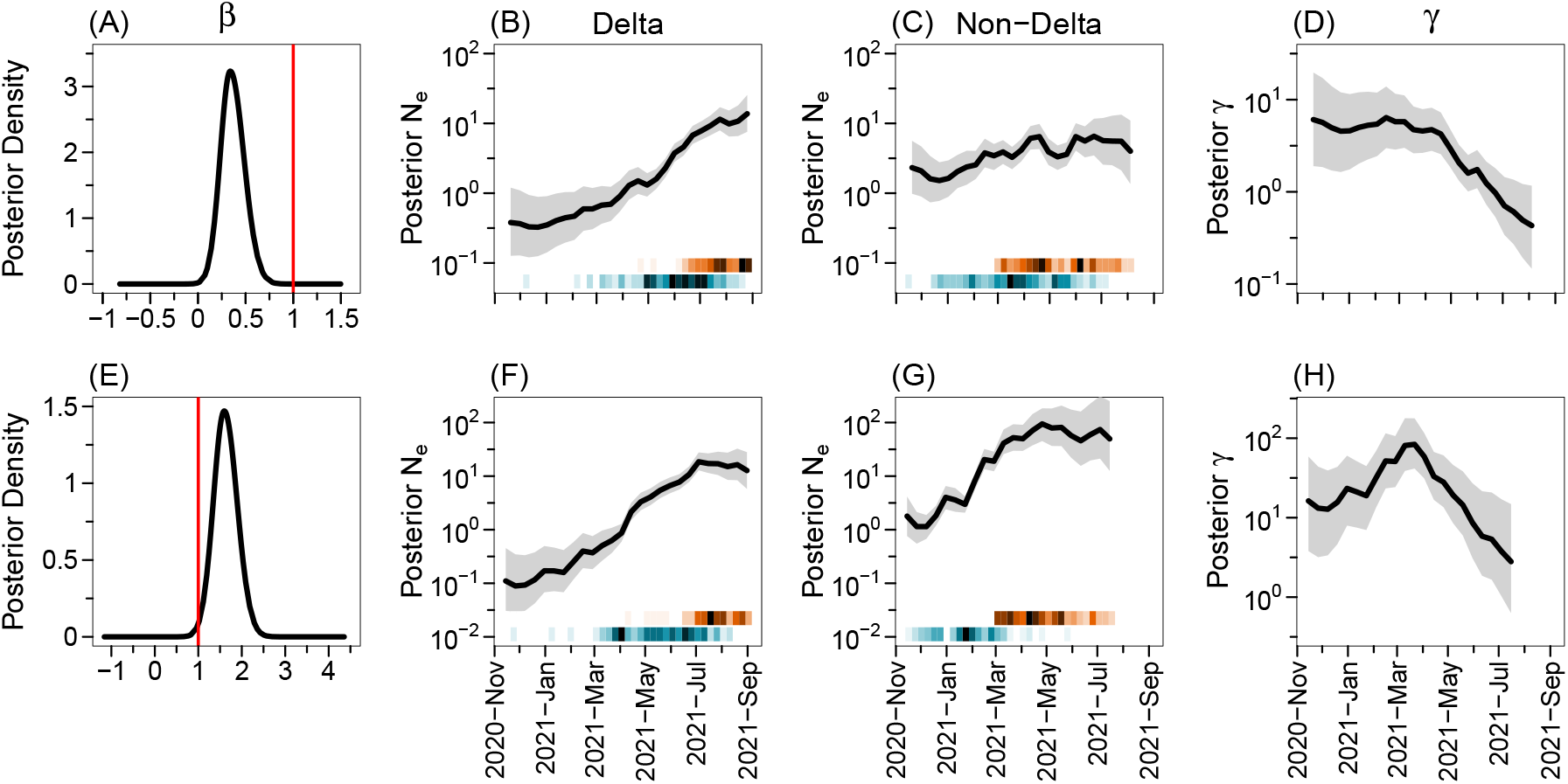
Posterior Inference of SARS-CoV-2 Population Dynamics in South Korea and Italy. Panels (A)-(D) contain results of South Korea, and the panels (E)-(H) shows results of Italy with **g^A^**=delta and **g^B^**=non-delta. The first column shows the *β* parameter posterior density from the parPop method. The red line indicates the value of *β* under the hypothesis that both variants share the same effective population size trajectory in the parPop model. The other columns present the results with adaPop+Pref: the second and the third columns display posterior estimates of EPS of the delta and non-delta variant, respectively, and the last column shows posterior estimates of *γ*. The solid line indicates the posterior medians with its surrounding shaded areas representing 95% BCIs. The orange and blue heatmaps describe the sampling and coalescent event intensity, respectively: the darker the color, the more number of events occurs in a time interval. The *y*-axis of plots in the columns 2-4 is plotted on a log scale. The results using other methods can be found in the Supplementary Materials.

Figure 4(B), (C), (F) and (G) depict the estimated posterior distribution of EPS of the two viral populations in the two countries obtained with adaPref+Pop (noPop estimates are qualitatively similar, see Supplementary Material). In both countries, the delta population has experienced prolonged growth since its inception, halting its growth in the last month. On the other hand, the non-delta population EPS grew until approximately the end of May (the growth looks more pronounced in Italy), then it roughly plateaued.

We next examine the quantification of the dependence between the two populations: *β* for parPop and *γ* for adaPop+Pref. In Figure 4(A), the posterior density of *β* estimated under the parPop model has mean 0.36 with 95% credible interval (0.19,0.51), well below 1 (red line), suggesting EPS growth is more pronounced among sequences having the delta variant in South Korea. On the other hand, the posterior density of *β* has mean 1.62 with credible interval (1.25,1.95) (Figure 4(E)) indicating the EPS of the non-delta variant grows faster than the delta variant in Italy under the parPop model. The result suggests that the growth of the non-delta population dominated that of the delta population which contradicts the general consensus (Mlcochova et al.,2021; del Rio et al., 2021).

Under the adaPop+Pref model, the monotonic decrease in the growth of the non-delta variant EPS compared to that of the delta variant EPS in South Korea is apparent in *γ*-trajectory (Figure 4(D)); this is consistent with parPop *β*. However, the *γ*-trajectory of Italy (Figure 4(H)) shows that, compared to the growth of the delta variant EPS, the non-delta variant underwent an initial phase of faster growth and then transitioned to slower growth around mid-March of 2021. The inability of the parPop model (Figure 4(E)) to capture the two-phase population dynamics in Italy, that are evident in the adaPop+Pref model (Figure 4(H)), suggests that more flexible approaches proposed by our work are needed for accommodating the broad range of population dynamics scenarios encountered in real applications.

## 5 Discussion

We have developed a coalescent-based Bayesian methodology for inferring dependent population size trajectories and quantifying such dependency. We make minimal assumptions on the functional form of population size trajectories and allow the dependence between the two populations to vary over time. We also present a sampling-aware model for leveraging additional information contained in sampling times for reduced bias and improved inference accuracy. Although the proposed models have an increased number of parameter, we prove that the models are identifiable. We have shown that our adaPop+Pref outperforms other methods in synthetic data with known ground truth and that our adaptive method can detect changes in population size dynamics that are otherwise undetected with other models. We make this point more precise in our SARS-CoV-2 analyses.

We have decided to infer parameters via a numerically approximated method that relies on Laplace approximations (INLA) for computational speed. However, our proposed models can, in principle, be implemented in any of the MCMC standard approaches. Implementing our models in an MCMC approach would allow to infer population size trajectories from molecular sequence and sequencing time information directly and it is a subject of future development. This extension would account for uncertainty in the genealogy, an important component that is missing in the analyses presented.

We see a growing number of applications of the coalescent requiring the modeling of complex demographic histories. In the introduction, we mentioned a few, such as viral epidemiology and cancer evolution. There is extensive literature on models incorporating more and more realistic features, for example, a detailed description of migration histories. The quest for realism and scientifically meaningful parameters comes at the cost of computational tractability and leads, sometimes, to issues of model identifiability. Our work is somewhat motivated by these problems. Our method is equipped with high performance and accuracy, due to its scalability, interpretability, and parameter identifiability; such properties are lacking in many complex models in biology and epidemiology. We see our proposal as a “hybrid approach” that allows scientists to quantify the relative advantage of one population over another while still retaining a fairly parsimonious model. Such approach will be invaluable across many biomedical disciplines for studying complex time-varying dependent evolutionary dynamics of populations.

