## Supplementary Materials for "Bayesian Inference of Dependent Population Dynamics in Coalescent Models"

### Bayesian Inference of Dependent Population Dynamics in Coalescent Models Supplementary Materials

#### 1 PROOFS OF THE RESULTS OF SECTION 3.4

We first recall a few preliminary definitions and results that will be used throughout the proofs. The review that follows is based on Lehmann & Casella (2006) Chapter 2.

Let $\mathbf{t}$ be distributed according to density $p_{\boldsymbol{\theta}}$ with respect to the Lebesgue measure, with $\boldsymbol{\theta} \in \Theta$, where $\boldsymbol{\theta}$ is a vector of length $s$. The *Expected Fisher Information* (EFI) is defined as $I(\boldsymbol{\theta}) = ||I_{i,j}(\boldsymbol{\theta})||$, where

$$I_{i,j}(\boldsymbol{\theta}) = E_{\boldsymbol{\theta}}\left[\frac{\partial \log p_{\boldsymbol{\theta}}}{\partial \theta_i}\frac{\partial \log p_{\boldsymbol{\theta}}}{\partial \theta_j}\right].$$

$I(\boldsymbol{\theta})$ depends on a given parametrization. Suppose that $\theta_i = h_i(\xi_1,\dots,\xi_p)$, for $i = 1,\dots,s$ and let $J$ denote the Jacobian matrix $J = ||\frac{\partial \theta_i}{\partial \xi_j}||$. Then the EFI $I(\boldsymbol{\xi})$ can be written as a function of $I(\boldsymbol{\theta})$ as follows

$$I(\boldsymbol{\xi}) = J\,I(\boldsymbol{\theta}(\boldsymbol{\xi}))\,J^T. \tag{S1}$$

If $p_{\boldsymbol{\theta}}$ belongs to an exponential family, then $\boldsymbol{\theta}$ is identifiable if $I(\boldsymbol{\theta})$ is nonsingular in a convex set containing $\Theta$ (Rothenberg (1971) Theorem 3). In Section 3 of the main text, we assumed that the EPS is piecewise-constant, $N_e = (N_{e,i})_{1:M}$, with the boundary points described by a regular grid $(k_i)_{1:M+1}$. Under the coalescent process, we have that $\mathbf{g}$ is distributed according to the coalescent density $p_{N_e}$ given in Eq. (1) in the main text, and $p_{N_e}$ belongs to the exponential family; i.e., to show that $N_e$ is identifiable, we need to show that $I(N_e)$ is nonsingular.

In the case of a single population, one can show that $I(N_e)$ is a diagonal matrix with diagonal entries $(c_1 N_{e,1}^{-2},\dots,c_M N_{e,M}^{-2})$, where $c_i$ denotes the number of coalescent times in the interval $[k_i, k_{i+1})$. The determinant $|I(N_e)|$ is non-zero if and only if $c_i > 0$ for all $i = 1\dots,M$. If this is the case, then $N_e$ is identifiable since $I(N_e)$ is full-rank, hence non-singular. The identifiability of $N_e$ is also studied by Parag & Pybus (2019).

The simplest model of two populations is to treat them independently: $\mathbf{g}^A$ is distributed to a coalescent with EPS $N_e^A = (N_{e,i}^A)_{1:M}$ and $\mathbf{g}^B$ is distributed to a coalescent with EPS $N_e^B = (N_{e,i}^B)_{1:M}$. For the same argument described above, $(N_e^A, N_e^B)$ are identifiabile, being $I(N_e^A, N_e^B)$ a $2M \times 2M$ diagonal matrix with entries $(c_1^A N_{e,1}^{A,-2}, c_1^B N_{e,1}^{B,-2},\dots,c_M^A N_{e,M}^{A,-2}, c_M^B N_{e,M}^{B,-2})$.

The hierarchical models described in Eqs. (4) and (5) can be seen as reparametrization of the independent model. To prove Propositions 1 and 2, we will use $I(N_e^A, N_e^B)$ and apply Eq. (S1). The two models define two different reparametrizations $\boldsymbol{\xi}$ and $J$ will be different.

##### 1.1 Proof of Proposition 1

*In model Eq. (4), we have that $N_e^A = (N_{e,i})_{1:M}$ and $N_e^B = (\gamma_i N_{e,i})_{1:M}$; hence, $\boldsymbol{\xi} = (N_{e,1},\dots,N_{e,M},\gamma_1,\dots,\gamma_M)$ and $J$ is a $2M \times 2M$ matrix. In this case, the Jacobian matrix is*

$$J = \begin{bmatrix} 1 & 1 & 0 & 0 & \dots \\ 0 & 1 & 0 & 0 & \dots \\ 0 & 0 & 1 & 1 & \dots \\ 0 & 0 & 0 & 1 & \dots \\ \vdots & \vdots & \vdots & \vdots & \ddots \end{bmatrix},$$

*hence it is a block diagonal matrix. The block diagonal structure follows from the independence of consecutive grid intervals. Its determinant is* 1 *being the product of the determinants of the subsquare* $2 \times 2$ *matrices along the diagonal. The result follows, because the determinant of the product of square matrices is the product of the determinants, i.e.,* $|I(N_e, \gamma)| \neq 0$ *and* $I(N_e, \gamma)$ *is nonsingular.*

##### 1.2 Proof of Proposition 2

*In model Eq. (5), we have that* $N_e^A = (N_{e,i})_{1:M}$ *and* $N_e^B = (\alpha N_{e,i}^{\beta})_{1:M}$*; hence,* $\boldsymbol{\xi} = (N_{e,1}, \ldots, N_{e,M}, \alpha, \beta)$ *and* $J$ *is a* $(M+2) \times 2M$ *matrix. We focus on the case* $M = 2$*. In this case, we have that the Jacobian matrix is*

$$J = \begin{bmatrix} 1 & \alpha\beta N_{e,1}^{\beta-1} & 0 & 0 \\ 0 & 0 & 1 & \alpha\beta N_{e,2}^{\beta-1} \\ 0 & N_{e,1}^{\beta} & 0 & N_{e,2}^{\beta} \\ 0 & \alpha N_{e,1}^{\beta} \log N_{e,1} & 0 & \alpha N_{e,2}^{\beta} \log N_{e,2} \end{bmatrix}.$$

*The determinant of* $J$ *is equal to* $N_{e,2}^{\beta} \alpha N_{e,1}^{\beta} \log N_{e,1} - N_{e,1}^{\beta} \alpha N_{e,2}^{\beta} \log N_{e,2}$*. This is different than zero because* $N_{e,1} \neq N_{e,2}$*. Hence,* $|J|$ *is non-zero. The result follows, because the determinant of the product of square matrices is the product of the determinants, i.e.* $|I(N_e, \alpha, \beta)| \neq 0$ *and* $I(N_e, \alpha, \beta)$ *is nonsingular.*

#### 2 SIMULATIONS DETAILS

*Details of the trajectories:* Here, we describe $N_e^A$ and $N_e^B$ for the six scenarios considered:

- Scenario 1:

$$N_e^A(t) = \begin{cases} 10 & \text{if } t \in [0, 0.35), \\ 10 \times 10^{14} \exp(-92.1034t) & \text{if } t \in [0.1, 0.3), \\ 0.1 & \text{if } t \in [0.4, \infty), \end{cases} \qquad N_e^B(t) = \frac{1}{2}(N_e^A(t))^{\frac{3}{2}}. \tag{S2}$$

- Scenario 2:

$$N_e^A(t) = \begin{cases} 10 & \text{if } t \in [0, 0.35), \\ 10 \times 10^{14} \exp(-92.1034t) & \text{if } t \in [0.1, 0.3), \\ 0.1 & \text{if } t \in [0.4, \infty), \end{cases} \qquad N_e^B(t) = \frac{1}{2} \quad \text{for all} \quad t. \tag{S3}$$

- Scenario 3:

$$N_e^A(t) = \begin{cases} 10 & \text{if } t \in [0, 0.35), \\ 10 \times 10^{14} \exp(-92.1034t) & \text{if } t \in [0.1, 0.3), \\ 0.1 & \text{if } t \in [0.4, \infty), \end{cases} \qquad N_e^B(t) = N_e^A(t + 0.1). \tag{S4}$$

- Scenario 4:

$$N_e^A(t) = \begin{cases} 10 & \text{if } t \in [0, 0.35), \\ 10 \times 10^{14} \exp(-92.1034t) & \text{if } t \in [0.1, 0.3), \\ 0.1 & \text{if } t \in [0.4, \infty), \end{cases} \qquad N_e^B(t) = N_e^A(t + 0.3). \tag{S5}$$

- Scenario 5:

$$N_e^A(t) = \begin{cases} \exp(13.04t - 4.61) & \text{if } t \in [0, 0.3), \\ 0.5 & \text{if } t \in [0.3, \infty), \end{cases} \qquad N_e^B(t) = \begin{cases} 4\exp(-12.29t) & \text{if } t \in [0, 0.3), \\ 0.1 & \text{if } t \in [0.3, \infty). \end{cases} \tag{S6}$$

- Scenario 6:

$$N_e^A(t) = \begin{cases} 10 & \text{if } t \in [0, 0.35), \\ 10 \times 10^{14} \exp(-92.1034t) & \text{if } t \in [0.1, 0.3), \\ 0.1 & \text{if } t \in [0.4, \infty), \end{cases} \qquad N_e^B(t) = N_e^A(t) \tag{S7}$$

*Simulation strategy:* We fix $n = 200$. For a given scenario, we set $n_1 = 2$, $s_1 = 0$, and simulate $n - n_1$ sampling locations from a Poisson process with rate 500. This defines $\mathbf{s}$ and $\mathbf{n} = (2, 1, \ldots, 1)$. We stress that the sampling is uniform, it does not depend neither on $N_e^A$ nor $N_e^B$, and the rate is chosen such that the expected number of samples in the interval $[0, 0.4]$ is equal to $n$. The interval $[0, 0.4]$ is of particular interest because it is where all the change points in $N_e^A$ and $N_e^B$ are concentrated. We repeat this step twice in order to define two pairs $(\mathbf{n}, \mathbf{s})$, which we label $(\mathbf{n}^A, \mathbf{s}^A)$ and $(\mathbf{n}^B, \mathbf{s}^B)$.

Conditionally on $\mathbf{n}^A$, $\mathbf{s}^A$, and $N_e^A$, we sample coalescent times $\mathbf{t}^A$ according to Algorithm 3 in Palacios & Minin (2013). We repeat the same step for $\mathbf{t}^B$. We sample from the inhomogeneous Poisson processes using the Lewis-Shedler thinning algorithm (Lewis & Shedler, 1979). Note that there is no need to sample tree topologies because the vector $\mathbf{t}$ is a sufficient statistic for $N_e(t)$.

### 3 GRID CONSTRUCTION

We rely on a discretization of $N_e$ and $\gamma$ defined by a regular grid $(k_i)_{1:M+1}$. A guideline on how to choose $M$ can be found in Faulkner et al. (2020). Throughout Section 3 of the main text, we assumed that $s_1^A$ and $s_1^B$ are equal to zero. In applications, this assumption might not hold. For example, the SARS-CoV-2 data sets in Section 4.2 where there are no samples of non-delta variant at time 0 (see heatmaps Figure 4).

Special care is required in this case. A condition for the identifiability of $(N_{e,1}, \ldots, N_{e,M}, \gamma_1, \ldots, \gamma_M)$ is that there should be at least one coalescent event per grid interval for both populations. Trivially, this condition will fail if one of the two populations does not have samples at the beginning of the grid.

A solution to this problem is the following. Use as population $A$, the population that has samples collected at time zero, and as $B$ the other population. Then, define the EPS of population $B$ as $N_e^B = (\gamma_i N_{e,i})_{L:M}$, where $L := \max_{i=1,\ldots,M}\{k_i \leq s_1^B\}$; i.e., truncate the effective population size in the intervals where there are no observations from $B$. The likelihood of $B$ does not contribute to $(N_{e,1}, \ldots, N_{e,L-1})$. In the analysis of SARS-CoV-2 data, we set the delta sample as population $A$, the non-delta as population $B$. Figure 4 depicts the truncation of $\gamma$: there are no estimates for $\gamma$ in the last few weeks.

### 4 SYNTHETIC DATA: ADDITIONAL RESULTS

Table S1 reports the average value of each statistic pooling together all scenarios and all grid points. We also average the performance metrics of $N_e^A$ and $N_e^B$. Table S1 differs from the table in the main manuscript because here, we are splitting the performance of each method by scenario.

adaPop+Pref stands out again as the best performing method. It only struggles to recover $N_e^B$ in Scenario 2: neither adaPop nor adaPop+Pref can recover the flat EPS as well as noPop and parPop. parPop is the best performing method in this case. The reason is that by simply setting $\beta = 0$, one can recover the flat $N_e^B$.

parPop is the method with the most polarized performance. When the parametric model is correctly specified (Scenarios 1, 2, and 6), it is among the best-performing methodologies. The accuracy deteriorates substantially when the model is misspecified (Scenarios 3, 4, and 5).

adaPop is often the second best performing method after adaPop+Pref. Surprisingly, it has a performance almost identical to parPop when parPop is correctly specified. This is surprising given that adaPop is more heavily parameterized. When parPop is misspecified, adaPop seems to adapt to the varying dependency and exhibit good performance.

### 5 SARS-CoV-2 MOLECULAR DATA ANALYSIS

For each country, we randomly sampled 150 sequences with delta variant and 150 sequences without delta variants collected in the period of 2021-03-01 to 2021-09-30 from publicly available high-coverage complete sequences in GISAID (Shu & McCauley, 2017). The date was chosen based on the availability of the first identified delta variants in each country (Figure S1). The resulting sampled sequence accession IDs can be found at the end of the Supplementary Materials. The classification of the delta variant group was based on the B.1.617.2 and AY lineages under the Pango lineage designation system (O'Toole et al., 2021). The sequences were aligned to the reference sequence EPI_ISL_402124 (Okada et al., 2020) with MAFFT (Katoh & Standley, 2013) by GISAID. From the multiple sequence alignment, we filtered out sites with more than 20% missing values in each country.

We analyzed sequences of each variant group of each country independently with BEAST2 (Bouckaert et al., 2019) using the Extended Bayesian Skyline prior on $N_e(t)$ (Heled & Drummond, 2008), the HKY mutation model with empirically estimated base frequencies (Hasegawa et al., 1985), and the strict clock model with the rate constrained to $[8 \times 10^{-4}, 1 \times 10^{-3}]$ substitutions per site per year (van Dorp et al., 2020). The chain was ran for $20 \times 10^6$ iterations, thinning every 1000 and with 20% burnin. The resulting maximum credibility clade (MCC) trees can be found in Figures S2 and S3.

With the inferred MCC trees, we examined the quantification of the dependence between the two variants in each country with four methods we introduced: noPop, parPop, adaPop, and adaPop+Pref (Section 4.1 in the main text). Results using the noPop, parPop, and adaPop methods are shown in Figures S4, S5, and S6, respectively, and the results with the adaPop+Pref method is shown in Figure 4 of the main text.

**Table S1.** Summary Statistics of Posterior Inference of $\gamma$, $N_e^A$, and $N_e^B$. Each entry is computed as the mean of the performance metric for a given scenario (100 datasets per scenario). The metrics for $N_e^A$ and $N_e^B$ have been also averaged. The numbers in bold indicate the method(s) with the best performance (and within 10% of the best) for each performance metric: the highest for ENV, the lowest for DEV and RWD.

| SCENARIO | METHOD | $\overline{\text{ENV}}\gamma$ | $\overline{\text{DEV}}\gamma$ | $\overline{\text{RWD}}\gamma$ | $\overline{\text{ENV}}N_e$ | $\overline{\text{DEV}}N_e$ | $\overline{\text{RWD}}N_e$ |
|---|---|---|---|---|---|---|---|
| 1 | adaPop+Pref | **1** | **0.26** | **2.74** | **0.98** | **0.3** | **2.53** |
| | adaPop | **1** | 0.3 | 4.33 | **0.98** | 0.47 | 6.74 |
| | noPop | **0.99** | 0.81 | 45.85 | **0.98** | 0.58 | 23.71 |
| | parPop | **1** | **0.24** | 9.7 | **0.97** | 0.46 | 7.05 |
| 2 | adaPop+Pref | **0.9** | **0.42** | 17.28 | **0.95** | 0.47 | 10.13 |
| | adaPop | **0.92** | 0.5 | 28.63 | **0.96** | 0.58 | 17.19 |
| | noPop | **0.93** | 0.62 | 469.22 | **0.97** | 0.28 | 3.32 |
| | parPop | **0.92** | **0.46** | 153.49 | **0.96** | **0.24** | **2.75** |
| 3 | adaPop+Pref | **0.9** | **0.35** | 2.12 | **0.91** | **0.27** | **1.65** |
| | adaPop | **0.94** | **0.36** | 3.2 | **0.94** | 0.32 | 2.67 |
| | noPop | **0.96** | 0.39 | 3.83 | **0.94** | **0.29** | 2.5 |
| | parPop | 0.4 | 1.16 | **1.86** | 0.52 | 0.77 | 2.05 |
| 4 | adaPop+Pref | 0.85 | **0.34** | **1.34** | **0.88** | **0.28** | **1.34** |
| | adaPop | **0.93** | **0.32** | 1.68 | **0.94** | 0.33 | 2.13 |
| | noPop | **0.96** | **0.34** | 2.1 | **0.93** | **0.26** | 1.86 |
| | parPop | 0.64 | 0.64 | 1.71 | 0.5 | 3.56 | 149.03 |
| 5 | adaPop+Pref | 0.71 | **0.52** | 1.55 | **0.88** | **0.27** | **1.48** |
| | adaPop | **0.8** | **0.52** | 2.07 | **0.92** | **0.28** | 1.81 |
| | noPop | **0.81** | **0.51** | 1.96 | **0.91** | **0.27** | 1.86 |
| | parPop | 0.52 | 0.64 | **1.35** | 0.73 | 0.8 | 1.56 |
| 6 | adaPop+Pref | **1** | **0.12** | **1.04** | **0.96** | **0.24** | **1.54** |
| | adaPop | **1** | **0.13** | 1.17 | **0.96** | 0.33 | 3.17 |
| | noPop | **0.99** | 0.47 | 9.14 | **0.96** | 0.4 | 5.66 |
| | parPop | **1** | **0.13** | 3.93 | **0.95** | 0.33 | 3.03 |

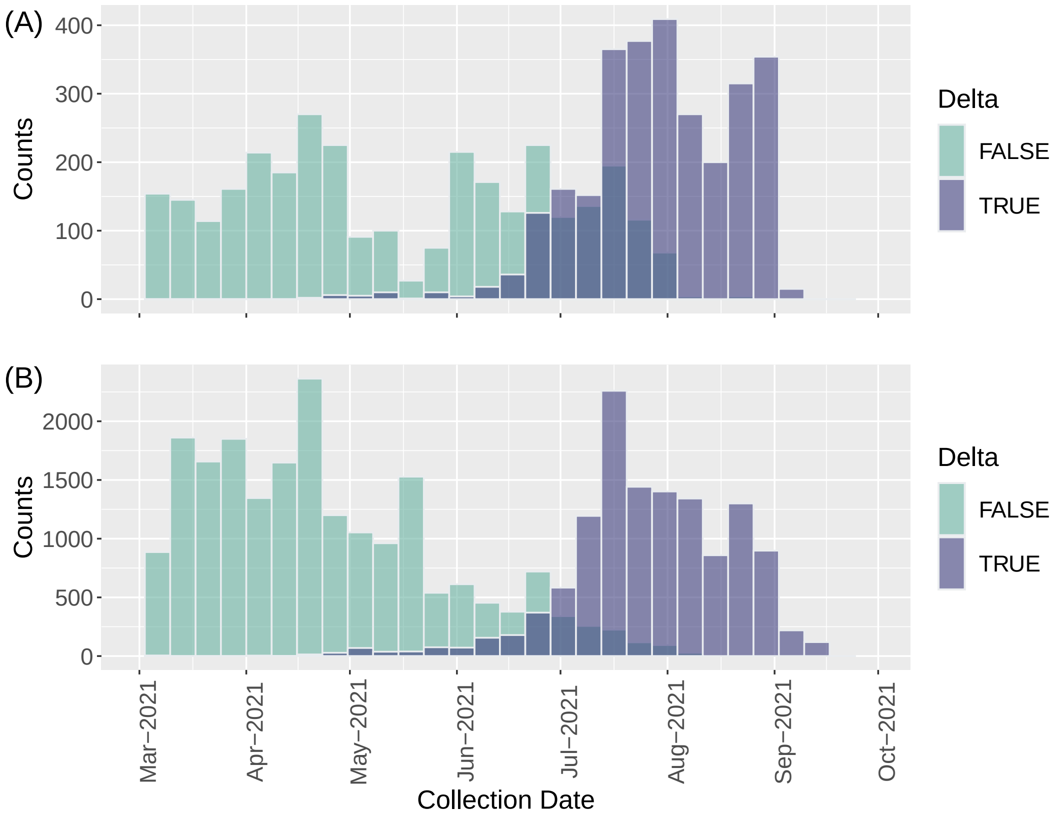


**Figure S1.** Collection Date Distributions of Available High-Coverage Complete Sequences in GISAID. (A) South Korea. (B) Italy. Purple and green colors indicate sequences with and without delta variants, respectively.

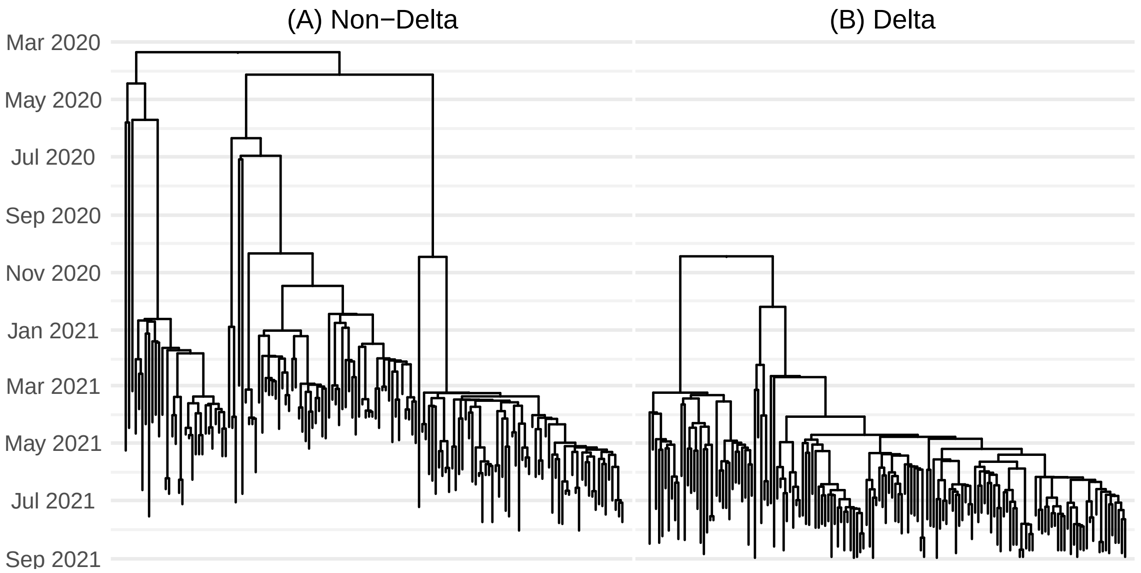


**Figure S2.** MCC Trees of Delta and Non-Delta Variants from South Korea.

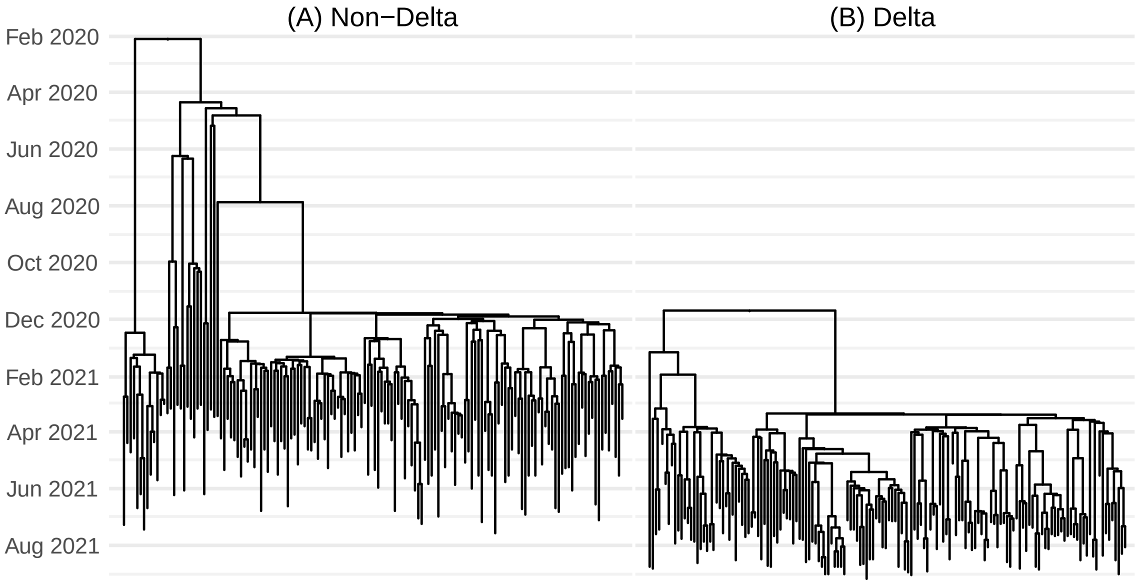


**Figure S3.** MCC Trees of Delta and Non-Delta Variants from Italy.

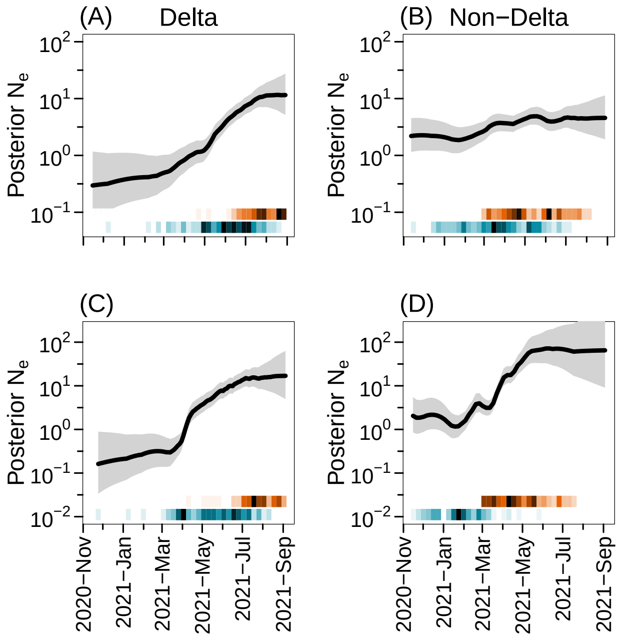


**Figure S4.** Posterior EPS Trajectories using the noPop Method. (A) South Korea, delta EPS. (B) South Korea, non-delta EPS. (C) Italy, delta EPS. (D) Italy, non-delta EPS. The figure format follows Figure 4 of the main text.

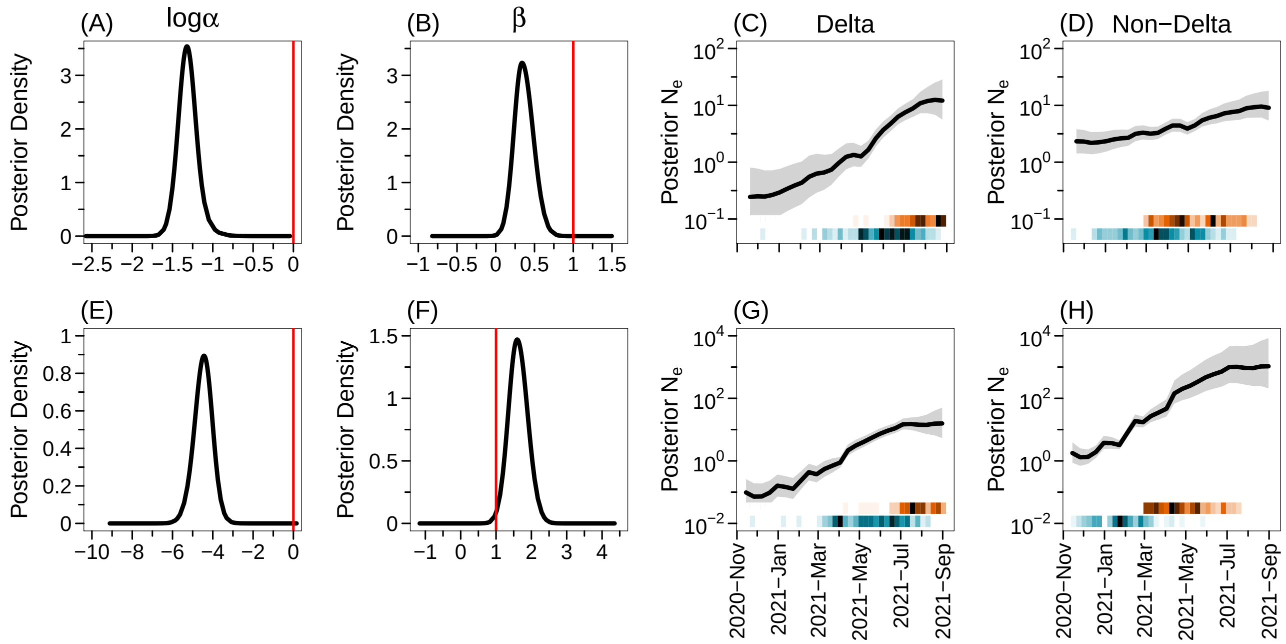

**Figure S5.** Posterior Densities of Parameters and Posterior EPS Trajectories using the parPop Method. (A) South Korea, $\log\alpha$. (B) South Korea, $\beta$. (C) South Korea, delta EPS. (D) South Korea, non-delta EPS. (E) Italy, $\log\alpha$. (F) Italy, $\beta$. (G) Italy, delta EPS. (H) Italy, non-delta EPS. The figure format follows Figure 4 of the main text.

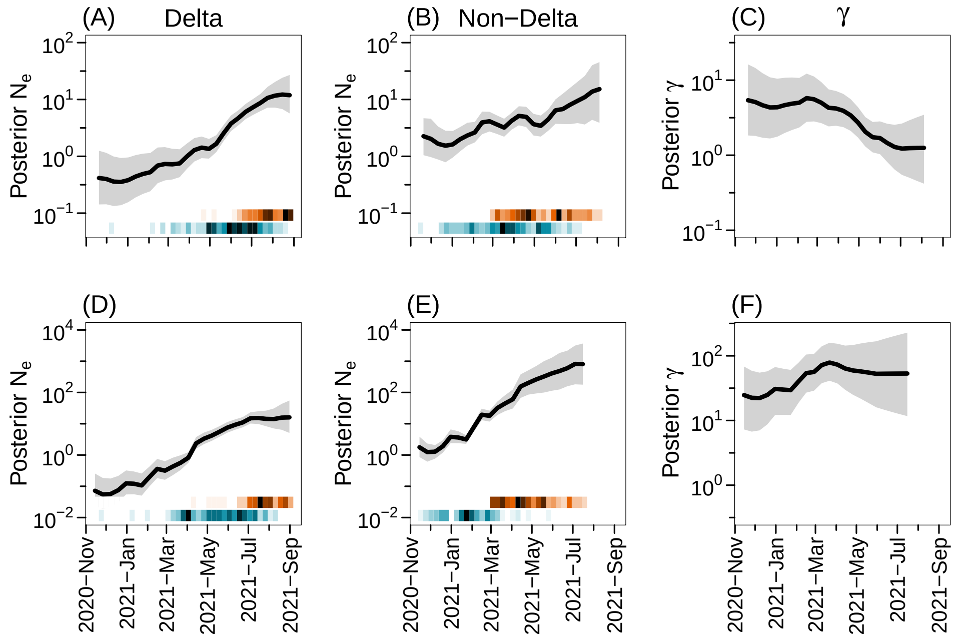

**Figure S6.** Posterior EPS Trajectories and Posterior Estimates of $\gamma$ using the adaPop Method. (A) South Korea, delta EPS. (B) South Korea, non-delta EPS. (C) South Korea, $\gamma$, (D) Italy, $\beta$. (E) Italy, delta EPS. (F) Italy, $\gamma$. The figure format follows Figure 4 of the main text.
